# Phylogenomic analyses of all major sea cucumber lineages provide a robust backbone of Holothuroidea

**DOI:** 10.1101/2025.04.14.648730

**Authors:** Jessica M. Whelpley, Abigail T. Uehling, Gustav Paulay, Joseph F. Ryan

## Abstract

Sea cucumbers (Echinodermata: Holothuroidea) are a diverse and ecologically important group found in benthic marine habitats across all latitudes and depths. We developed a phylogeny of sea cucumbers by applying transcriptome and genome data from 37 Holothuroidea species representing 7 orders and 13 families, including 16 transcriptomic datasets generated as part of this study, with a matrix of 4,900 translated transcripts encompassing 2,689,407 amino acid sites. We confirm the placement of a Apodida as sister to the remaining Holothuroidea. We were able to bring resolution to Neoholothuriida by recovering Persiculida as sister to Molpadida and Synallactida as sister to Dendrochirotida. We recovered four families (Stichopodidae, Synallactidae, Sclerodactylidae, and Cucumariidae) as polyphyletic. Our study highlights the need for formally revising existing taxonomic classifications of Holothuroidea and provides a foundation from which the evolution of this group of animals can be investigated.

## Introduction

The approximately 1,800 described species of sea cucumbers (Echinodermata: Holothuroidea; retrieved in August 2023 from World Register of Marine Species) are found across benthic environments from the equator to the poles, and from littoral to hadal depths (O’Loughlin et al. 2011). Species diversity is highest in the tropics while in some regions of the deep sea, sea cucumbers can represent >90% of animal biomass, making them one of the dominant “large” animals on our planet (Kerr & Kim 2001). While holothuroids are relatively conservative in their overall body plan, they have evolved some remarkable evolutionary innovations, such as evisceration, respiratory trees, and anal suspension feeding. Holothuroids are unusual among echinoderms in many aspects, including their worm-like, bilaterally symmetrical body plan and decreased skeletonization. In addition, they include some of the smallest and longest exemplars within Echinodermata, ranging from 2mm to > 3m (Miller et al. 2017). The group includes numerous epifaunal and burrowing clades, as well as a few groups capable of swimming as adults, including the only known pelagic echinoderm, *Pelagothuria natatrix* Théel, 1882. Sea cucumbers play an important ecological role as deposit feeding bioturbators.

Until recently, broad analyses of holothuroid relationships have relied on relatively few morphological traits (e.g., Haeckel 1896; Ludwig 1891; Perrier 1902); Smirnov (2012) provided an in-depth historical overview of the relationships based on morphology. Kerr & Kim (2001) carried out the first formal phylogenetic analysis of the class, based on 47 morphological characters. They recovered the Apodida, lacking tube feet and respiratory trees, to be sister to all other Holothuroidea, and the deep-sea Elasipodida as sister to the remaining lineages that are characterized by the possession of respiratory trees. These results, with minor adjustments, have held up to later phylogenetic scrutiny. In an overview relying on morphological characters but lacking a phylogenetic analysis, Smirnov (2012) made important adjustments to the classification, including the proposal that the lungless family Deimatidae, traditionally classified in the Elasipodida, is more closely related to sea cucumbers with respiratory trees and has lost these secondarily, and in beginning the breakup of the Synallactidae by separating the Mesothuriidae.

Lacey et al. (2005) published the first molecular tree based on 18S ribosomal RNA gene sequences from 16 species, which supported the relationships of the Apodida, Elasipodida, and lunged sea cucumbers, and recovered the order Aspidochirotida as paraphyletic.

Miller et al (2017) in the broadest phylogenetic analysis of holothuroids to date, with 82 species and 6 genes (3 mitochondrial, 3 nuclear), confirmed these results, including the placement of Deimatidae among the lunged sea cucumbers suggested by Smirnov (2012). They also subdivided the former Aspidochirotida into the Persiculida, Synallactida, and Holothuriida, erected two new families, and discussed further examples of family-level non-monophyly in the class.

### Objectives

Here we present a broadly-sampled phylogenomic analysis of Holothuroidea to test and refine sea cucumber relationships. Our phylogenomic analysis used data from two sources: 1) new transcriptome data for 16 species selected to fill current gaps, and 2) publicly available transcriptomes and genomes. Based on analyses of these data, we present a robust hypothesis of higher-level relationships in Holothuroidea that contributes to our understanding of sea cucumber biodiversity.

## Methods

### Reproducibility and Transparency Statement

Custom scripts, command lines, and data used in these analyses, including alignment and tree files, are available at https://github.com/josephryan/SeaCucumberPhylogenomics. To maximize transparency and minimize confirmation bias, analyses were planned *a priori* using a phylotocol (DeBiasse and Ryan 2018) and posted to our GitHub repository (URL above). A version of this GitHub Repo was archived at the time of submission and is available through the following DOI: 10.5281/zenodo.8253475.

### Specimen acquisition

Specimens were collected from shore, by SCUBA, remotely operated vehicles (ROVs), or purchased from aquarium stores, photographed live, and subsampled for tissues.

### RNA extraction and sequencing

Tissues were dissected from narcotized animals, rinsed in filtered sea water, placed in RNA*later*, and held in a refrigerator for 24 hours to allow buffer saturation of tissue, then stored at -80C. Total RNA extraction and library construction were performed at the University of Florida Interdisciplinary Center for Biotechnology Research (ICBR). Samples with a total RNA <100 ng were discarded. RNA-Seq sequencing of 150bp paired-end reads was performed with Illumina HiSeq 3000 also by ICBR.

### Transcriptome de novo assembly and quality control

We generated transcriptomic data from 16 sea cucumbers representing 6 orders and 11 families. Next, we identified and downloaded publicly available holothuroid transcriptome and genome datasets from various sources (Supplementary Table 1). We then used these data to construct a dataset of 37 holothuroid species representing 13 families from the 7 orders recognized by Miller et al. (2017) plus a single representative of each non-holothuroid echinoderm class. We used BUSCO v5 (Simao et al. 2015) to assess the completeness of each of these datasets. BUSCO completeness scores varied widely among holothuroid datasets with “complete” scores ranging from 5.97 - 99.48% (“complete + partial” scores ranged from 20.55 - 99.58%) with an average of 61.17% (“complete + partial” average scores were 77.30%; Supplementary Table 1). The average completeness score for the new transcriptomes was 65.60% (average “complete + partial” score for new transcriptomes was 81.73%).We selected outgroup datasets from multiple datasets that we assembled based on highest BUSCO scores.

We trimmed all RNA-Seq reads with BL-Filter (Dunn et al. 2013) and then assembled reads in Trinity v2.8.5 (Grabherr et al. 2011) with the “include_supertranscripts” parameter set to generate super-transcripts (Davidson et al. 2017). We used BUSCO v2/v3 (Benchmarking universal single-copy orthologs) through the gVolante platform (Nishimura et al. 2017; https://gvolante.riken.jp) to assess completeness of all transcriptomes. We used TransDecoder v3.0.1 (Haas & Papanicolaou 2016) to translate RNA sequences into amino acid sequences. We used Alien Index v2.1 (Ryan 2014; https://github.com/josephryan/alien_index) with the alien_index database v0.02 to identify and remove any sequences that had better BLAST hits to non-animal sequences than they did to animal sequences.

### Ortholog identification, alignment, phylogenetic analysis

To identify orthologous contigs among transcriptomes, we used diamond v0.9.22.123 (Buchfink et al. 2015) to perform reciprocal best BLAST searches on the 41 datasets, then generated groups of orthologous sequences (orthogroups) in OrthoFinder v2.2.3 (Emms and Kelly 2015).

We used MAFFT v7.309 (Yamada et al. 2016) to align protein sequences within orthogroups. We filtered paralogs from each transcriptome using PhyloPyPruner version 1.2.4 (Kocot et al. 2013; https://gitlab.com/fethalen/phylopypruner). The resulting alignments that possessed fewer than 50% sequence gaps were concatenated in a super matrix using fasta2phylomatrix version 0.02 (Ryan 2019; github.com/josephryan/JFR-PerlModules). We used IQ-Tree v1.5.5 (Nguyen et al. 2015) to estimate a maximum-likelihood (ML) phylogeny using modelfinder (built into IQ-Tree) for model assignments for each partition. This analysis included 1,000 ultrafast bootstrap replicates.

## Results

We collected and sequenced RNA from 16 species of sea cucumber (Figure 1). From these data and publicly available transcriptomes, we generated a concatenated alignment of 2,689,407 amino acid columns from 42 species with a 32% occupancy. Our phylogenomic analyses resulted in concordant topologies with 100% bootstrap support at each node (Figure 2; Supplementary Figure 1).

**Figure 1.**
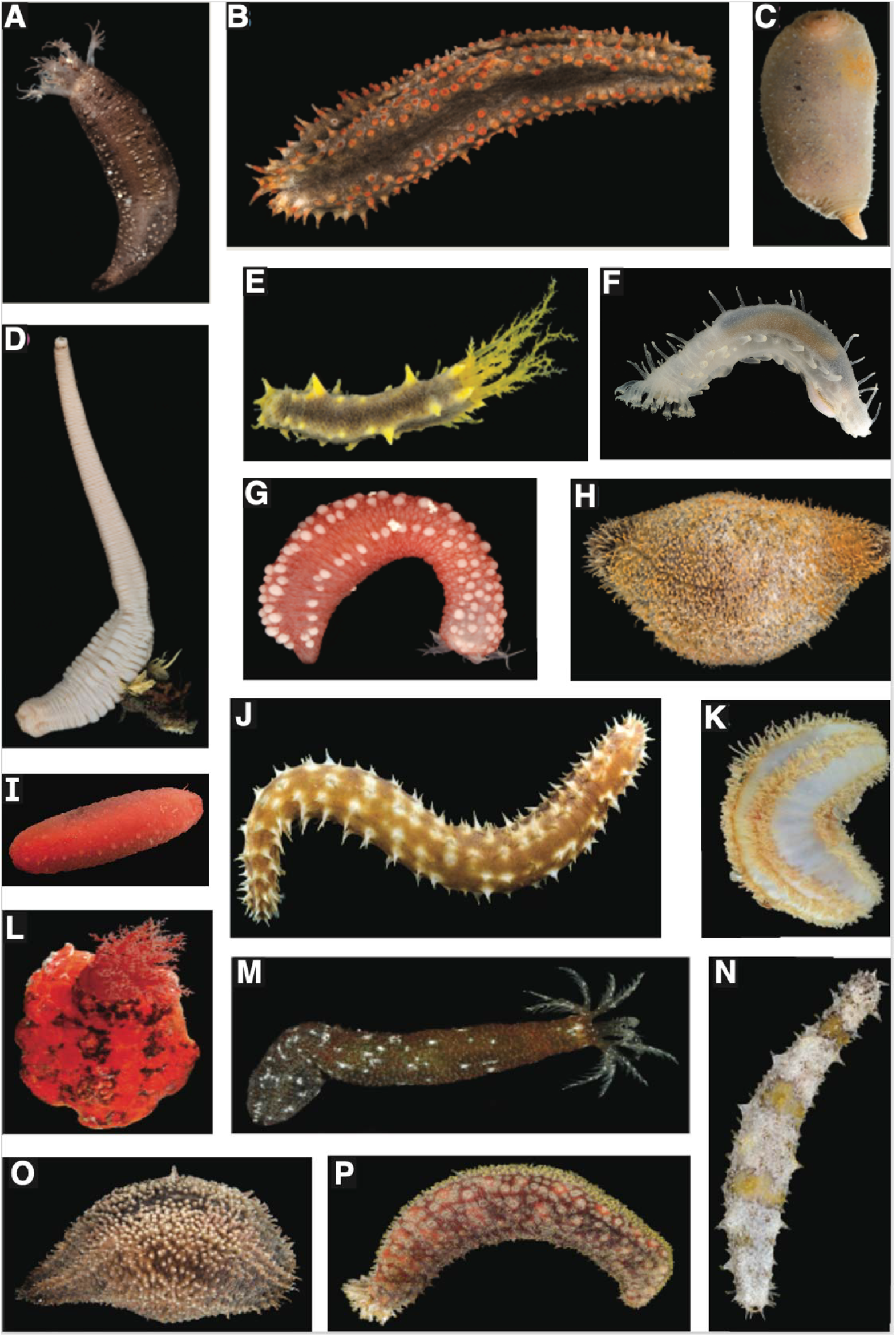
Representative holothuroid specimens included in this study. **A**. *Polycheira rufescens*, **B**. *Aslia pygmaea*, **C**. *Molpadia intermedia*, **D**. *Paracaudina chilensis*, **E**. *Colochirus robustus*, **F**. *Pannychia* sp., **G**. *Chiridota rigida*, **H**. *Sclerodactyla briareus*, **I**. *Synallactidae* sp., **J**. *Holothuria hilla*, **K**. *Pentamera pediparva*, **L**. *Psolus chitonoides*, **M**. *Synaptula hydriformis*, **N**. *Holothuria floridana*, **O**. *Thyonella* sp., **P**. *Holothuria grisea*. Images by G. Paulay.

**Figure 2.**
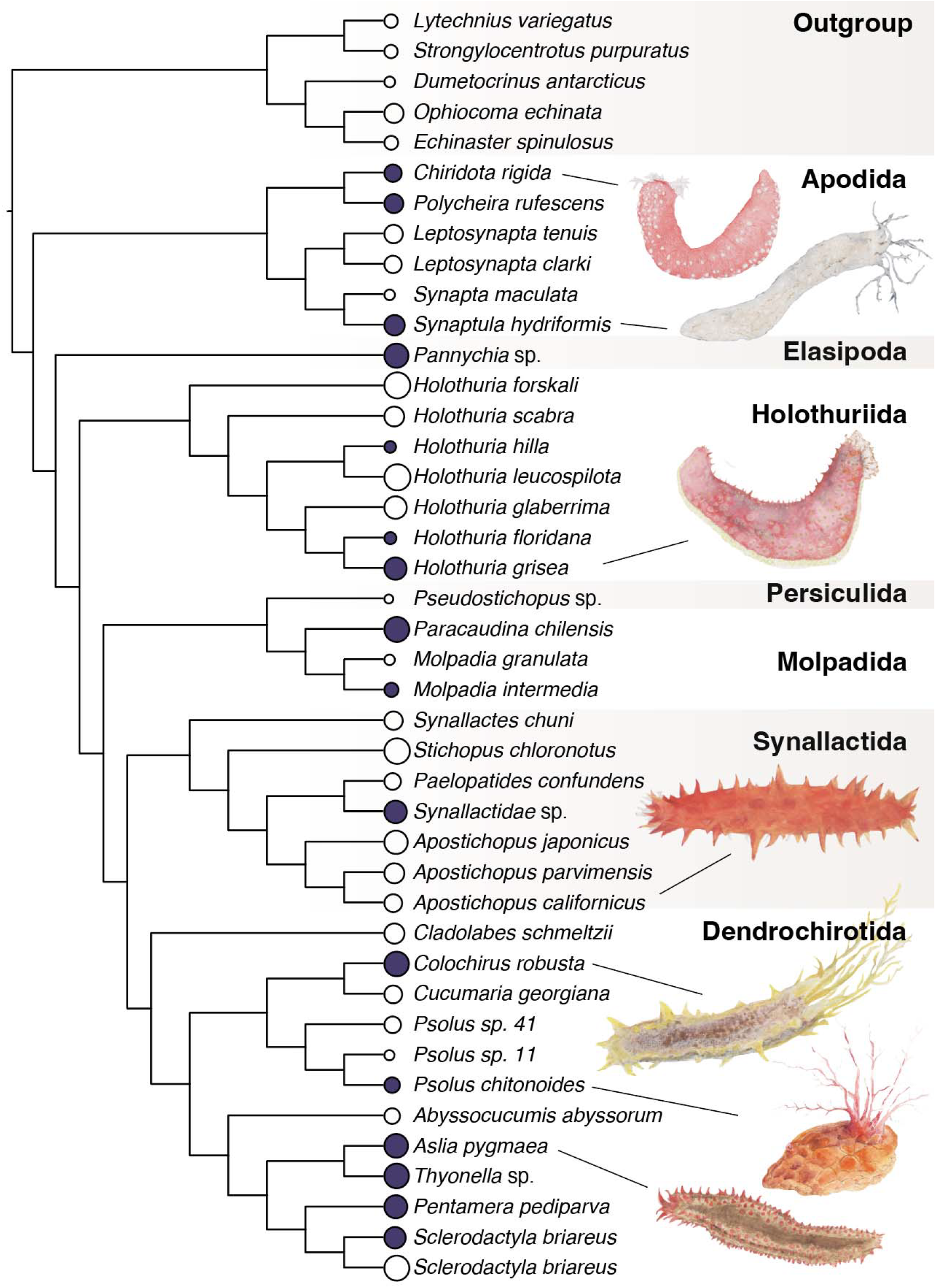
Concatenated maximum-likelihood phylogeny of Holothuroidea. All nodes have 100% support from ultrafast bootstraps. Circles at tips indicate matrix occupancy, with filled-in circles denoting sequence data new to this study and clear circles denoting previously published transcriptome or genome sequences. See Supplementary Figure 1 for version with branch lengths. Watercolors by M. Lara-Whelpley.

We recovered all seven Holothuroidea orders as monophyletic with all nodes fully resolved and with 100% bootstrap support for each node (Figure 2). We recovered a monophyletic Apodida as sister to Actinopoda, the remaining Holothuroidea. Elasipodida was sister to the Pneumonophora, within which, Holothuriida was sister to the Neoholothuriida, all consistent with previous results (Miller et al. 2017). Our inferred phylogeny resolves the previously uncertain relationships within the Neoholothuriida, with Persiculida sister to Molpadida, and Synallactida sister to Dendrochirotida.

Of the 9 families represented by multiple taxa, four were polyphyletic: Sclerodactylidae, Stichopodidae, Synallactidae, and Cucumariidae. Within the Cucumariidae, the subfamilies Cucumariinae and Colochirinae were also polyphyletic. The five genera that included multiple species were all monophyletic. Taxonomic assignments of studied species are included in Supplemental Table 2.

## Discussion

Holothuroidea is a diverse clade of echinoderms dating to the Middle Ordivician (464 million years ago; Reich 2010). Understanding the origin and evolution of this group requires a robust understanding of the higher-level relationships in the clade. Many hypotheses of the relationships have been posited (e.g., Kerr and Kim 2001; Lacey et al. 2005; Miller et al. 2017, Mongiardino Koch et al. 2023). Here we present a firmly resolved holothurian phylogeny with transcriptomic and genomic data consisting of representatives from all 7 orders and 13 of the recognized families (Figure 2).

The relationship of sea cucumbers was recently overhauled in a 6-gene phylogeny of 82 species, the first in-depth phylogenetic assessment of the class (Miller et al. 2017). The results indicated the traditional order Aspidochirotida and several families are polyphyletic, recognized three new orders, two new families, and outlined the broad evolutionary history of the group. We find support for most clades erected by this study and further clarify relationships of some. We recovered all seven orders defined by Miller et al. (2017) as monophyletic, albeit two were represented by single species. Relationships among orders were consistent with their results, except for relationships within the Neoholothuriida that were weakly supported in that study.

Several morphological characters have received attention as informative for the deep relationships of sea cucumbers, especially the presence of podia and respiratory trees and the morphology of tentacles and skeletal elements. Miller et al. (2017) provided robust support for deep divisions among sea cucumbers that lack podia and respiratory trees (Apodida), those that possess podia but not respiratory trees (Elasipodida), and all others that have both (or have secondarily lost one). These divisions and taxa have been discussed since the late 1800’s (Ludwig 1889-1892, Haeckel 1896; see review by Smirnov 2012) and phylogenetically assessed by Kerr & Kim (2001).

### Apodida

Apodida (sea cucumbers without tube feet) consist of three families Myriotrochidae, Chiridotidae and Synaptidae and have the earliest representation in the fossil record for crown-group holothuroids in the early/middle Silurian (Reich, 2010). Our results support the deep division between the Apodida and all other holothuroids that have tube feet (Figure 2). Ludwig (1889-1892) named the latter Actinopoda, a name resurrected by Miller et al. (2017).

The apodid families Chiridotidae and Synaptidae and subfamilies Leptosynaptinae and Synaptinae sampled were monophyletic in our study (Figure 2). Previous work recovered these two families as non-monophyletic with modest support (Miller et al. 2017). The two species underlying this conflict in Miller et al. (2017), *Leptosynapta clarki* and *Chiridota rigida*, were both represented in our analyses. It will be important for future phylogenomic studies to include a representative from the Myriotrochidae, considered the sister to the other apodid families based on morphological and genetic data (Smirnov 2012; Miller et al. 2017).

### Actinopoda (Elasipodida, Holothuriida, Persiculida, Molpadida, Synallactida, Dendrochirotida)

Actinopoda (sea cucumbers with tube feet) consist of the six orders and are divided into Elasipodida (sea cucumbers without respiratory trees) and Pneumonophora (sea cucumbers with respiratory trees). We recovered Elasipodida (sea cucumbers without respiratory trees) as the sister group to the rest of Actinopoda (Figure 2) as in previous studies (Kerr & Kim 2001; Miller et al. 2017).

### Pneumonophora (Holothuriida, Persiculida, Molpadida, Synallactida, Dendrochirotida)

As in recent studies, we recovered Pneumonophora (sea cucumbers with respiratory trees). Furthermore, our results support the separation of Synallactidae into the Persiculida and Synallactidae.

### Holothuriida

Within Pneumophora, we recovered Holothuriida (sea cucumbers with gonads on one side of the dorsal mesentery) as sister to the remaining clades (Figure 2), consistent with Miller et al. (2017). Holothuriida consists of two families, Holothuriidae and Mesothuriidae. Our analyses include seven species of Holothuriidae, a group which includes some of the best known sea cucumbers. A defining synapymorphy of Holothuriidae are their sticky cuvierian tubules that are digestive diverticula used as defense. Numerous Holothuriidae species are consumed, and some in our study, including *Holothuria glaberrima* and *Holothuria scabra*, are model organisms for developmental research (Medina-Feliciano et al. 2021; Hamel et al. 2001). Most holothuriids are in the genus *Holothuria* which is subdivided into numerous subgenera based mostly on endoskeletal characters. Unfortunately, our study lacks representatives of Mesothuriidae.

### Neoholothuriida (Persiculida, Molpadida, Synallactida, Dendrochirotida)

Miller et al. (2017) erected Neoholothuriida as the sister lineage to the remaining Pneumonophora, however, the relationship of its four lineages remained ambiguous due to low support values. Our analyss, which greatly expand the number of genes applied to this question, provide resolution for this clade with strong support. Within Neoholothuriida, we recovered the Persiculida as sister to Molpadida and the Synallactida as sister to Dendrochirotida (Figure 2).

### Persiculida

Persiculida (a clade characterized by sparse or no body wall ossicles) was erected based on modest branch support, mostly for species previously assigned to the Synallactidae (Miller et al. 2017). While our study includes only one persiculid, its robust placement as sister to molpadids supports the polyphyly of Synallactidae and recognition of Persiculida. This relationship makes sense of the unstable position of the enigmatic genus *Gephyrothuria*, which has been placed in either Synallactidae or Molpadiida based on morphological evidence (O'Loughlin 1998), and found to fall in the Persiculida based on sequence data (Miller et al. 2017).

### Molpadida

Molpadida (sea cucumbers without tube feet but with respiratory trees) were at times grouped with Apodida because they lack podia (e.g., Brandt 1835; Perrier 1902), but this relationship has not been supported by modern analyses (e.g., Miller et al. 2017). Our results support the placement of Molpadida as sister to Persiculida.

### Synallactida

Synallactida was erected for the restricted Synallactidae and the related Stichopodidae. The latter includes some of the most important sea cucumbers in fisheries and aquaculture. These two families were intermixed, not monophyletic, as found by Miller et al. (2017). *Synallactes* and *Stichopus*, the type genera of the two families, were sister to other genera. The Synallactidae is differentiated from the Stichopodidae by the absence of tentacle ampullae.

### Dendrochirotida

Dendrochirotida (sea cucumbers with an introvert and retractor muscles) is the most diverse group of sea cucumbers. Most species in this group are suspension feeders with highly branched, dendritic tentacles. Our analyses recovered Dendrochirotida as monophyletic and included 4 (Sclerodactylidae, Cucumariidae, Phyllophoridae, Psolidae) of the 13 families within the group. Both Cucumariidae (5 species) and Sclerodactylidae (2 species) were polyphyletic in our analyses as was the case in Miller et al.’s (2017) study, which found Psolidae and Phyllophoridae to be polyphyletic as well, a relationships that we were unable to test because of limited taxon sampling.

## Conclusions

Using 4,900 protein coding genes, we present a robust hypothesis of higher-level relationships in Holothuroidea that contributes to our understanding of sea cucumber evolution. Our findings confirm the support for most clades erected by Miller et al. (2017) in a study based on far fewer characters. Our results support the findings of Apodida, a tube-feet lacking sea cucumber, as sister to the rest of Holothuroidea. Contrary to Miller et al. (2017) we recovered the two sampled apodid families (Chiridotidae and Synaptidae) as monophyletic, supporting the morphological characters uniting them. This study further confirms that Elasipodida is a deeply diverging lineage that is sister to the rest of Actinopoda. We recovered Holothuriida as sister to the Neoholothurida, consistent with Miller et al. (2017). Low support values in previous work left open the question as to the evolutionary relationships of the remaining lineages in Neoholothurida. Our analyses recovered Persiculida as sister to Molpadida and Synallactida sister to Dendrochirotida with full support in all our analyses. As in previous studies, we found several families to be polyphyletic. These results suggest the need for taxonomic revisions and highlight the need for additional taxon sampling. Furthermore, the work highlights several clades that provide powerful phylogenetic frameworks from which to investigate the gain and loss of complex morphological and physiological traits.

## Additional

### Funding

Funding for JW was provided by a University of Florida Biodiversity Institute (UFBI) Fellowship and sequencing was provided by a UFBI SEED Fund Awarded to JFR and GP.

## Acknowledgements

We would like to thank Steven Haddock for inviting GP on a cruise and the captain and crew of the R/V Western Flyer and the pilots of the ROV Doc Ricketts of Monterey Bay Aquarium Research Institute, who enabled the collection of deep-sea specimens for this study. We would also like to thank Daniel A. Janies and Greg E. Rouse for providing transcriptome sequences from their U.S. National Science Foundation Assembling the Echinoderm Tree of Life project [grant numbers DEB 1036219, 1036368]. Additionally, we would like to thank the staff at the Interdisciplinary Center for Biotechnology Research Gene Expression Core (RRID:SCR_019135) and NextGen Sequencing Core (RRID:SCR_019152) for their work. We thank Pamela S. Soltis and Matthew E. Smith for their feedback. JW would especially like to thank Rebecca M. Varney for her feedback, insight and support.

## Supporting Information

GitHub Repository with phylotocol: DOI: 10.5281/zenodo.8253476 NCBI/ENA BioProject: PRJEB73365

