## Supplementary Table 1 for "Phylogenomic analyses of all major sea cucumber lineages provide a robust backbone of Holothuroidea"

**Supplemental** **Table 1.** Taxonomic information and SRA accession numbers for specimens used in the molecular analyses. Results of BUSCO assessment for completeness and gene occupancy of concatenated gene matrices are included. Classes are denoted by letters: H = Holothuroidea, A = Asteroidea, C = Crinoidea, E = Echinoidea, O = Ophiuroidea. Specimen ID: voucher catalog numbers in UF Echinodermata

| **Class** | **Species** | **SRA** | **Specimen ID** | **BUSCO**  **(c / c+p) %** | **Matrix Occupancy %** |
| --- | --- | --- | --- | --- | --- |
| H | *Abyssocucumis abyssorum* | SRR2830762 | NA | 27.67 / 54.51 | 53.14 |
| H | *Apostichopus californicus* | SRR1139198 | NA | 41.40 / 71.28 | 60.45 |
| H | *Apostichopus japonicus* | ASM275485v1 | NA | 84.38 / 93.19 | 79.22 |
| H | *Apostichopus parvimensis* | SRR2484238 | NA | 99.48 / 99.58 | 67.84 |
| H | *Aslia pygmaea* | PRJEB73365 | 18052 | 87.11 / 95.81 | 80.29 |
| H | *Chiridota rigida* | PRJEB73365 | NA | 44.34 / 72.33 | 58.82 |
| H | *Cladolabes schmeltzii* | SRR6023958 | NA | 57.86 / 73.17 | 68.22 |
| H | *Colochirus robustus* | PRJEB73365 | 18053 | 90.57 / 95.91 | 83.41 |
| H | *Cucumaria georgiana* | SRR3190098 | NA | 45.07 / 67.61 | 60.47 |
| H | *Holothuria floridana* | PRJEB73365 | 18056 | 19.08 / 39.20 | 40.82 |
| H | *Holothuria forskali* | SRR5109955 | NA | 92.98 / 96.23 | 88.08 |
| H | *Holothuria glaberrima* | SRR490864 | NA | 69.50 / 89.31 | 76.14 |
| H | *Holothuria grisea* | PRJEB73365 | 18054 | 84.80 / 94.13 | 75.67 |
| H | *Holothuria hilla* | PRJEB73365 | NA | 18.24 / 40.25 | 39.69 |
| H | *Holothuria leucospilota* | DRR023763 | NA | 96.23 / 98.53 | 88.65 |
| H | *Holothuria scabra* | SRR5755244 | NA | 51.68 / 75.26 | 67.35 |
| H | *Leptosynapta clarki* | SRR1695478 | NA | 58.39 / 82.60 | 58.92 |
| H | *Leptosynapta tenuis* | SRR3217898 | NA | 82.29 / 91.72 | 62.63 |
| H | *Molpadia granulata* | SRR2845419 | NA | 5.97 / 21.17 | 35.45 |
| H | *Molpadia intermedia* | PRJEB73365 | NA | 36.90 / 59.64 | 48.00 |
| H | *Paelopatides confundens* | BJ42 | NA | 36.90 / 64.26 | 56.14 |
| H | *Pannychia* sp*.* | PRJEB73365 | NA | 87.53 / 95.39 | 81.82 |
| H | *Paracaudina chilensis* | PRJEB73365 | NA | 86.90 / 94.13 | 84.65 |
| H | *Pentamera pediparva* | PRJEB73365 | NA | 71.70 / 88.47 | 78.00 |
| H | *Polycheira rufescens* | PRJEB73365 | 19331 | 78.30 / 90.25 | 64.31 |
| H | *Psolus* sp. 11 | BJ11 | NA | 69.08 / 86.90 | 34.20 |
| H | *Psolus* sp. 41 | BJ41 | NA | 30.19 / 62.05 | 56.33 |
| H | *Psolus chitonoides* | PRJEB73365 | NA | 53.88 / 71.07 | 52.82 |
| H | *Pseudostichopus* sp. | BJ44 | NA | 8.49 / 20.55 | 30.12 |
| H | *Sclerodactyla briareus* | SRR1139189 | NA | 94.65 / 97.48 | 84.69 |
| H | *Sclerodactyla briareus* | PRJEB73365 | 18051 | 57.86 / 75.89 | 72.45 |
| H | *Stichopus chloronotus* | SRR2846098 | NA | 86.48 / 94.65 | 86.65 |
| H | *Synallactes chuni* | SRR2895367 | NA | 33.54 / 68.24 | 62.31 |
| H | Synallactidae sp*.* | PRJEB73365 | NA | 71.28 / 87.21 | 76.44 |
| H | *Synapta maculata* | SRR2846103 | NA | 20.65 / 37.74 | 35.78 |
| H | *Synaptula hydriformis* | PRJEB73365 | 18057 | 91.19 / 97.48 | 69.20 |
| H | *Thyonella* sp. | PRJEB73365 | 18055 | 88.78 / 95.70 | 83.47 |
| A | *Echinaster spinulosus* | [SRR1139455](https://trace.ncbi.nlm.nih.gov/Traces/sra/?run=SRR1139455) | NA | 98.43 / 99.06 | 46.88 |
| C | *Dumetocrinus antarcticus* | SRR5564112 | NA | 96.65 / 98.74 | 36.31 |
| E | *Lytechinus variegatus* | [SRR1139214](https://trace.ncbi.nlm.nih.gov/Traces/sra/?run=SRR1139214) | NA | 94.03 / 98.22 | 45.59 |
| E | *Strongylocentrotus purpuratus* | SAMN00829422 | NA | 99.37 / 99.69 | 45.18 |
| O | *Ophiocoma echinata* | [SRR1138707](https://trace.ncbi.nlm.nih.gov/Traces/sra/?run=SRR1138707) | NA | 76.73 / 89.52 | 65.12 |

**Supplemental** **Table 2.** Taxonomic assignment of studied species.

| **Class** | **Order** | **Family** | **Species** |
| --- | --- | --- | --- |
| Holothuroidea | Dendrochirotida | Cucumariidae | *Abyssocucumis abyssorum* (Théel, 1886) |
| Holothuroidea | Synallactida | Stichopodidae | *Apostichopus californicus* (Stimpson, 1857) |
| Holothuroidea | Synallactida | Stichopodidae | *Apostichopus japonicus* (Selenka, 1867) |
| Holothuroidea | Synallactida | Stichopodidae | *Apostichopus parvimensis* (H.L. Clark, 1913) |
| Holothuroidea | Dendrochirotida | Cucumariidae | *Aslia pygmaea* (Théel, 1886) |
| Holothuroidea | Apodida | Chiridotidae | *Chiridota rigida* (Semper, 1867) |
| Holothuroidea | Dendrochirotida | Sclerodactylidae | *Cladolabes schmeltzii* (Ludwig, 1875) |
| Holothuroidea | Dendrochirotida | Cucumariidae | *Colochirus robustus* (Östergren, 1898) |
| Holothuroidea | Dendrochirotida | Cucumariidae | *Cucumaria georgiana* (Lampert, 1886) |
| Holothuroidea | Holothuriida | Holothuriidae | *Holothuria floridana* (Pourtalès, 1851) |
| Holothuroidea | Holothuriida | Holothuriidae | *Holothuria forskali* (Delle Chiaje, 1824) |
| Holothuroidea | Holothuriida | Holothuriidae | *Holothuria glaberrima* (Selenka, 1867) |
| Holothuroidea | Holothuriida | Holothuriidae | *Holothuria grisea (*Selenka, 1867 |
| Holothuroidea | Holothuriida | Holothuriidae | *Holothuria hilla* Lesson, 1830 |
| Holothuroidea | Holothuriida | Holothuriidae | *Holothuria leucospilota* (Brandt, 1835) |
| Holothuroidea | Holothuriida | Holothuriidae | *Holothuria scabra* Jaeger, 1833 |
| Holothuroidea | Apodida | Synaptidae | *Leptosynapta clarki* Heding, 1928 |
| Holothuroidea | Apodida | Synaptidae | *Leptosynapta tenuis* (Ayres, 1851) |
| Holothuroidea | Molpadida | Molpadiidae | *Molpadia granulata* (Ludwig, 1893) |
| Holothuroidea | Molpadida | Molpadiidae | *Molpadia intermedia* (Ludwig, 1893) |
| Holothuroidea | Synallactida | Synallactidae | *Paelopatides confundens* (Ludwig, 1893) |
| Holothuroidea | Elasipodida | Laetmognidae | *Pannychia* sp*.* |
| Holothuroidea | Molpadida | Caudinidae | *Paracaudina chilensis* (J. Müller, 1850) |
| Holothuroidea | Dendrochirotida | Phyllophoridae | *Pentamera pediparva* Lambert, 1998 |
| Holothuroidea | Apodida | Chiridotidae | *Polycheira rufescens* (Brandt, 1835) |
| Holothuroidea | Persiculida | Pseudostichopodidae | *Pseudostichopus* sp. |
| Holothuroidea | Dendrochirotida | Psolidae | *Psolus* sp. 11 |
| Holothuroidea | Dendrochirotida | Psolidae | *Psolus* sp. 41 |
| Holothuroidea | Dendrochirotida | Psolidae | *Psolus chitonoides* Clark, 1901 |
| Holothuroidea | Dendrochirotida | Sclerodactylidae | *Sclerodactyla briareus* (Lesueur, 1824) |
| Holothuroidea | Synallactida | Synallactidae | *Stichopus chloronotus* Brandt, 1835 |
| Holothuroidea | Synallactida | Synallactidae | *Synallactes chuni* Augustin, 1908 |
| Holothuroidea | Synallactida | Synallactidae | *Synallactida* sp*.* |
| Holothuroidea | Apodida | Synaptidae | *Synapta maculata* (Chamisso & Eysenhardt, 1821) |
| Holothuroidea | Apodida | Synaptidae | *Synaptula hydriformis* (Lesueur, 1824) |
| Holothuroidea | Dendrochirotida | Cucumariidae | *Thyonella* sp. |

**Supplemental Figure 1.** ML phylogeny of Holothuroidea with branch lengths. All nodes have 100% bootstrap support. Circles at tips indicate matrix occupancy, with filled-in circles denoting sequence data new to this study and clear circles denoting previously published transcriptome or genome sequences.

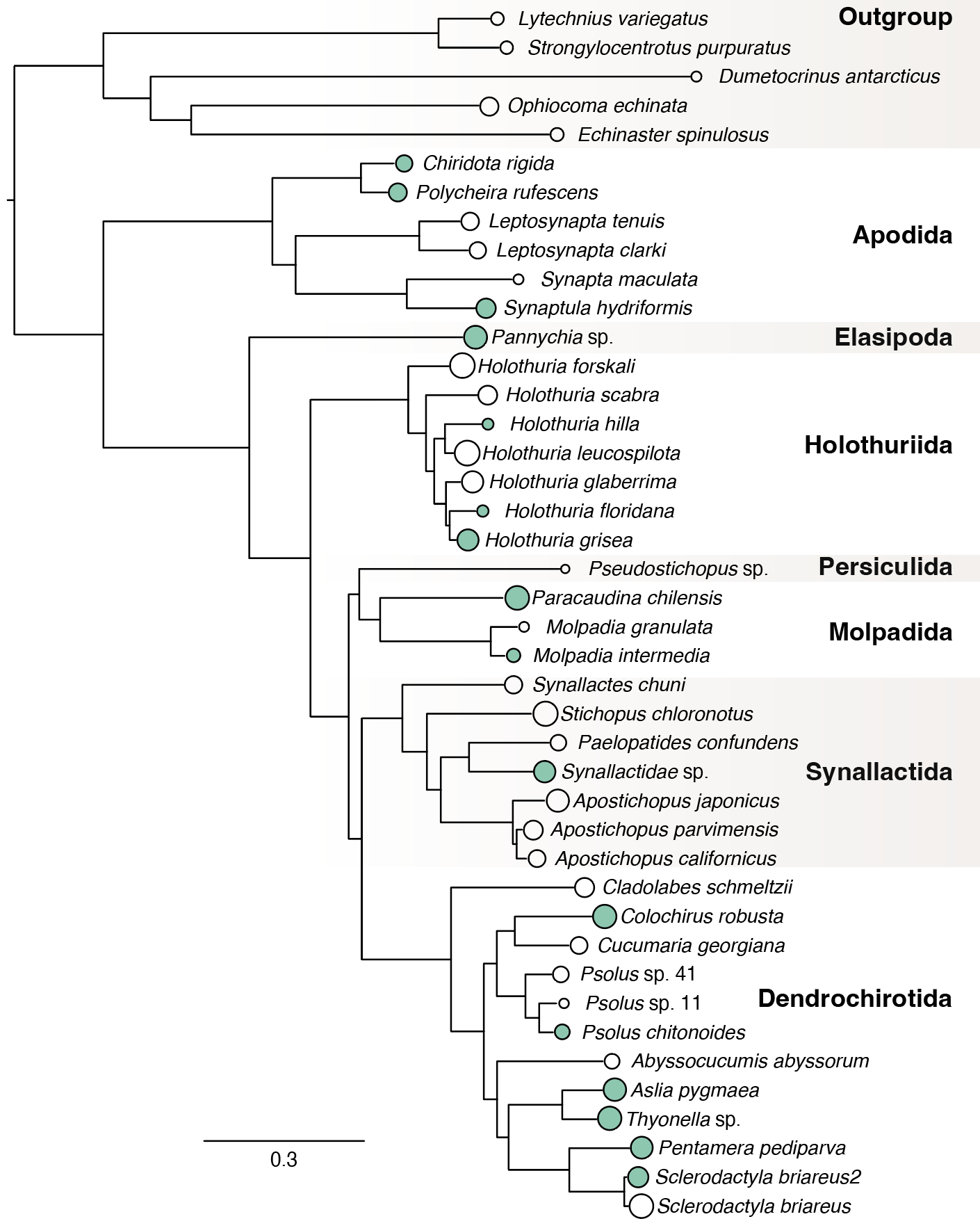
